# Action information contributes to metacognitive decision-making

**DOI:** 10.1101/657957

**Authors:** Martijn E. Wokke, Dalila Achoui, Axel Cleeremans

**Author notes:** These authors contributed equally to this work (Martijn E. Wokke, Dalila Achoui).

## Abstract

Monitoring and control of our decision process are key ingredients of adept decision-making. Such metacognitive abilities allow us to adjust ongoing behavior and modify future decisions in the absence of external feedback. Although metacognition is critical in many daily life settings, it remains unclear what information is actually being monitored and what kind of information is being used for metacognitive decisions. In the present study, we investigated whether response information connected to perceptual events contribute to metacognitive decision-making. Therefore, we recorded EEG signals during a perceptual color discrimination task while participants were asked to provide an estimate about the quality of their decision on each trial. Critically, the moment participants provided second-order decisions varied across conditions, thereby changing the amount of action information (e.g., response competition or response fluency) available for metacognitive decisions.

Results from three experiments demonstrate that metacognitive performance improved when first-order action information was available at the moment metacognitive decisions about the perceptual task had to be provided. This behavioral effect was accompanied by enhanced functional connectivity (beta phase synchrony) between motor areas and prefrontal regions, exclusively observed during metacognitive decision-making. Our findings demonstrate that action information contributes to metacognitive decision-making, thereby painting a picture of metacognition as a second-order process, integrating sensory evidence and the state of the decider during decision-making.

**Significance:** Monitoring and control of our decision process is a critical part of every day decision-making. When feedback is not available, metacognitive skills enable us to modify current behavior and adapt prospective decision-making. Here, we investigated what kind information is being used to compute an estimate about the quality of our decisions. Results demonstrate that during perceptual decision-making, information about one’s actions towards perceptual events is being used to evaluate the quality of one’s decisions. EEG results indicate that functional connectivity between motor regions and prefrontal cortex could serve as a mechanism to convey action information during metacognitive decision-making. Considered together, our results demonstrate that post-decisional information contributes to metacognition, thereby evaluating not only what one perceives (e.g., strength of perceptual evidence) but also how one responds towards perceptual events.

## Introduction

The ability to monitor and evaluate the quality of our decision-making is crucial for adept behavior. For instance, when driving a car for a long time it is important to have a reliable estimate about the adequacy of ones driving performance to avoid unsafe situations. However, not much is known how our brain constructs such an estimate, or what exactly is being monitored and evaluated. In lab settings, perceptual or memory tasks have been frequently used to probe the mechanisms that underpin metacognitive performance (Morales et al. 2018). In such studies, first-order task performance generally correlates with second-order (metacognitive) decisions, leading to the intuitive assumption that metacognitive decisions are largely based on the same information that governs first-order decision-making (Kiani and Shadlen 2009; Yueng and Summerfield 2012; Fetsch et al. 2014).

In recent years, however, dissociations between objective task performance and subjective ratings, and dissociations between sources of information supporting first- and second-order decisions have been observed (Wierzchoń et al. 2014; Fleming et al. 2015; Berg et al. 2016; Wokke et al. 2017; Palser et al. 2018). Typically, metacognitive decisions follow first-order responses, thereby allowing certain sources of information to become available during second-order decision-making. Recent findings suggest that metacognition can be supported by ‘embodied’ processes, such as interoception or response information that become available for metacognitive decision-making after a first-order decision has been made (Cisek and Kalaska 2005; Wierzchoń et al. 2014; Fleming et al. 2015; Allen et al. 2016; Urai et al. 2017; Palser at al. 2018). For instance, manipulation of neural activity via transcranial magnetic stimulation over premotor cortex resulted in altered confidence judgments during a perceptual task (Fleming et al. 2015). Critically, stimulation of premotor areas reduced metacognitive capacity without changing visual discrimination performance. Further, it has been shown that the order of rating confidence (before or after the response) influenced metacognitive performance on an anagram problem-solving task (Siedlecka et al. 2016). From a computational perspective, Pasquali and colleagues explored neural network architectures aimed at capturing the complex relationships between first-order and second-order (metacognitive) performance in a range of different cognitive tasks and suggested that metacognitive judgments are rooted in learned redescriptions of first-order error information rather than in the relevant first-order information itself (Pasquali et al. 2010). This is broadly consistent with Fleming and Daw’s perspective, in which they offered to unify the above observations in a single framework in which confidence operates as a second-order computation about one’s own performance (Fleming and Daw 2017). In this framework, samples of sensory evidence that support first- and second-order decisions are coupled yet distinct. Interestingly, their second-order model of confidence computation incorporates knowledge about the reliability of actions towards perceptual events.

Here, in three experiments, we aimed to elucidate whether and in what way action information informs metacognitive judgments. We therefore constructed a color discrimination task in which we varied the amount of available action information (i.e., response strength and fluency of response execution) at the moment a metacognitive judgment had to be provided. Our design enabled us to contrast metacognitive decisions based on purely perceptual information (uninformed by action processes) with metacognitive decisions having access to both perceptual and motor action information. We recorded electroencephalographic signals to investigate whether functional connectivity between motor regions and prefrontal cortex could serve as a mechanism to convey relevant action information (e.g., response competition or response fluency) during metacognitive decision-making.

Previously, beta oscillations have been intimately linked to sensory and motor processing (Pfurtscheller and Lopes da Silva 1999). Recently, however, beta-band power (de)synchronization in motor regions has been shown to provide insight into the dynamics underlying perceptual decisions (Donner et al. 2009) and response uncertainty (Tzagarakis et al. 2010). Beta oscillations have repeatedly been shown to predict first-order decisions (Donner et al. 2007; Donner et al. 2009; Haegens et al. 2011), to support maintenance of persistent activity (Siegel et al. 2012; Engel et al. 2010; Kloosterman et al. 2015) to mediate long-range communication, and to play an important role in the preservation and ‘awakening’ of endogenous information (Spitzer and Haegens, 2017). Here, we focused on beta phase synchrony between motor regions and prefrontal cortex (Wokke et al. 2017). Specifically, we expected both functional connectivity (beta phase synchrony) and metacognitive performance to increase when response information about first-order decisions would be accessible during metacognitive decision-making.

## Materials and Methods

### Participants

Twenty-five participants (15 females, mean age= 21.1, SD= 4.82) took part in experiment 1, twenty-nine participants (18 females, mean age= 22.1, SD= 2.65) in experiment 2, and twenty (13 females, mean age= 21.6, SD= 3.87) in the control experiment. Participants received financial compensation for their participation in this experiment. All participants had normal or corrected-to-normal vision and were naïve to the purpose of the experiment. All procedures complied with international laws and institutional guidelines and were approved by the local Ethics Committee, and all participants provided their written informed consent prior to the experiment.

### Task design

A field of 600 green and red moving dots was centrally presented (250*250 pixels) on a Dell 17 monitor with a refresh rate of 60 Hz. The monitor was placed at a distance of ∼57 cm in front of each participant so that the collection of moving dots subtended a visual angle of 6.6°. On each trial, a field of green and red colored randomly moving dots were presented (3*3 pixels in diameter) against a black background. Crucially, on each trial a majority of the 600 dots (on average 315.11 dots, SD=6.76) was either green or red. Participants were instructed to determine what color (red or green) was predominant on each trial by pressing a left (∼) or right (/) key. The level of difficulty was determined for each participant individually by using a one-up-two-down staircase procedure in steps of 0.5% of total number of dots^83^ before the start of the experiment. After two consecutive correct responses, the difference between the total number of red vs. green dots was reduced by 0.5%. During the staircase procedure, each participant performed a total of three blocks (one block of each condition in experiment 1, and one block of each condition in the control and second experiment plus a block randomly picked between condition 1 and 2) in order to assess the level of difficulty that resulted in a stable level of performance set at 75% correct. The stimulus was presented for 800 ms, and at any moment during stimulus presentation a total of 600 dots were displayed. Each trial started with a blank screen (jittered between 1000-1500 ms, in steps of 100 ms) on which a fixation cross was centrally presented. After stimulus presentation a blank was presented for 1000 ms to avoid the influence on prolonged evidence accumulation (Yueng and Summerfield 2012; Hebart et al. 2014).

### Experiment 1

In the first experiment, we created three conditions by varying the amount of available action information at the moment a metacognitive decision had to be provided (Figure 1a). In condition 1, the stimulus and blank screen were followed by a response cue (2000 ms), instructing participants whether the left or right button corresponded to the answer “green” or “red” (Figure 1a). The stimulus-response mapping was randomized so that in approximately half of all trials the left response button signaled ‘red’ and in approximately the other half of the trials it signaled ‘green’. This randomized stimulus-response mapping prevented participants from preparing their response immediately after the visual stimulus had appeared and enabled us to disentangle motor preparation from motor action in both our behavioral and EEG analyses. After the presentation of the response cue, participants were asked to indicate whether the majority of the dots were green or red by pressing the corresponding button with their left or right index finger. Next, participants had to provide a metacognitive judgment about their decision by indicating their level of confidence in being correct on a labeled scale from 1-4, where 1 indicated being very uncertain and 4 being very certain that their first-order response was correct. Participants were encouraged to use the whole range of the scale. Participants verbally reported their confidence rating in order to link the manual motor response exclusively to the first-order decision (red-green decision). A microphone registered all verbal responses using speech recognition software in Presentation (Neurobehavioral Systems, version 18.1), allowing automatic recording of verbal responses. To ensure an accurate transcription of the responses, we set a threshold level of certainty (0.8). Flagged trials below 0.8 certainty were checked manually and corrected if necessary (4% of all trials).

**Figure 1.**
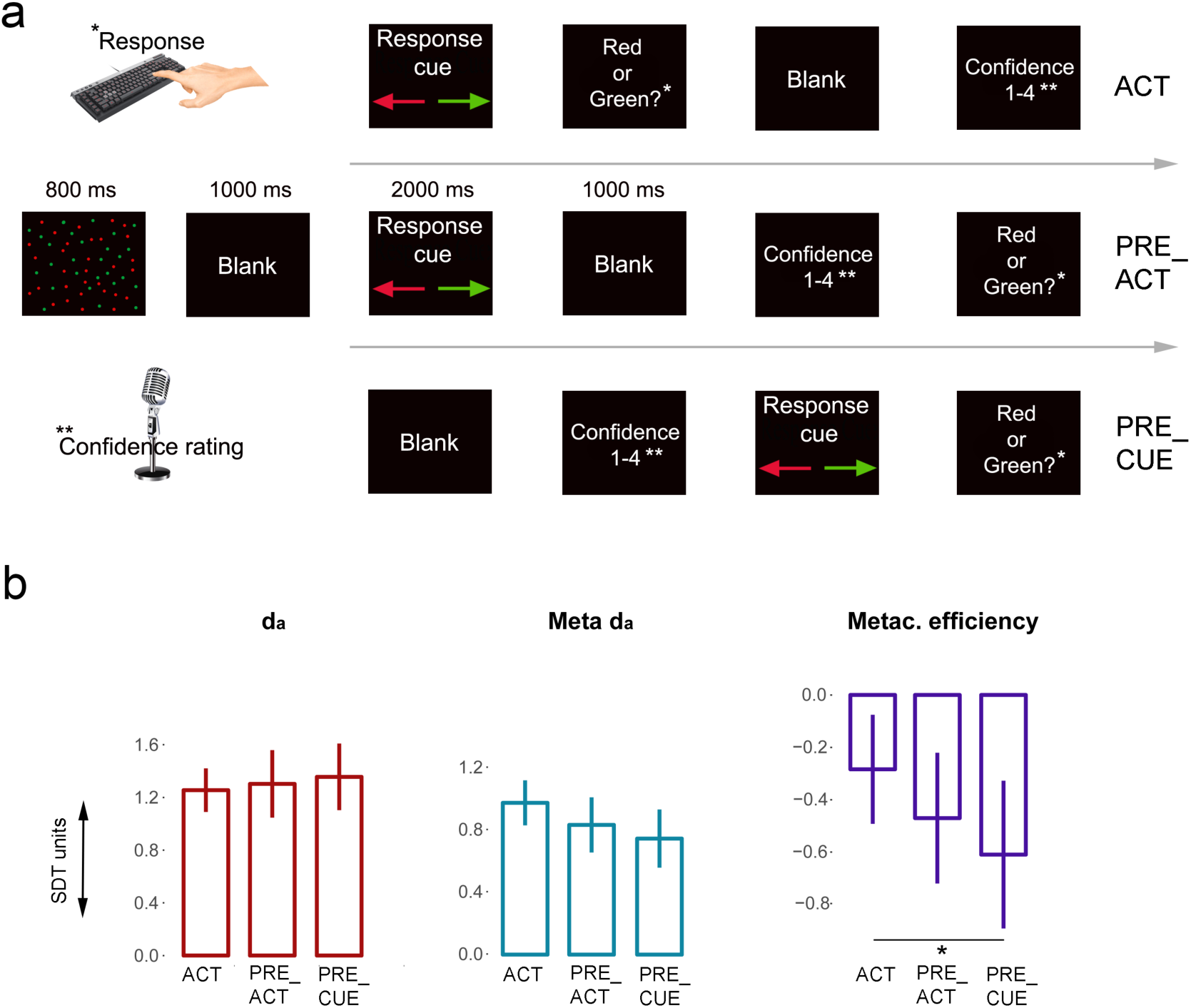
a) *Task design experiment 1.* Participants had to decide whether the majority of randomly moving dots were red or green by pressing a left or right key. The key that mapped onto a ‘red’ or ‘green’ answer was signaled by a response cue on each trial. Verbal confidence ratings were recorded either at the end of each trial (ACT), or directly preceding the first-order response (PRE_ACT), or directly following stimulus presentation (PRE_CUE). In this way, in each condition a different amount of first-order action information was available at the moment metacognitive decisions were provided. B) *Behavioral results.* Participants’ metacognitive efficiency decreased when action information was not available, while first-order performance remained unaltered. Error bars represent within-subjects standard error of the mean.

Critically, confidence ratings were given at different points in the trial sequence depending on the condition. Typically, confidence ratings are given after the first-order task response (ACT). However, in this experiment we manipulated the amount of action information (i.e., response execution, action preparation) available for metacognitive decisions by varying the position of metacognitive judgments in a trial. In PRE_ACT, metacognitive judgments had to be provided before the first-order response (after the response cue), while in PRE_CUE metacognitive decisions had to be made prior to the first-order response and presentation of the response cue. This resulted in two conditions in which action information was minimal (response preparation) or absent at the moment the second-order (metacognitive) decision was made.

### Control experiment

In the control experiment, we investigated whether observed EEG results were specific to metacognitive processes, by studying the non-specific effect of epiphenomenal/lingering motor activity from first-order responses. Therefore, we used a similar task design as used in the first experiment. Critically, in the control experiment participants were instructed to verbally report one randomly chosen letter out of four presented letters (‘E, ‘G’, ‘P’, ‘T’), instead of providing a confidence rating. Here, we focused on differences between ACT and PRE_ACT, since we did not observe behavioral and functional connectivity differences between PRE_ACT and PRE_CUE in the first experiment, see Figure 1b.

### Experiment 2

In the second experiment, the response cue was removed in order to establish reliable stimulus-response mappings throughout the experiment. The rest of the design was kept similar to that of the first experiment (Figure 1c).

### Behavioral analyses

In the present experiment, we aimed to investigate whether we could observe changes in metacognitive (second-order) performance depending on condition. In order to obtain a reliable measure of changes in second-order performance we used a staircase procedure before starting the experiment (see above) and filtered the data by excluding participants whose d_a_ or meta d_a_ scores were <0.5 or >2.0 in ACT of the three experiments (the position of the metacognitive judgment in ACT is typically used when measuring metacognitive performance). For analyses, 15 participants were included in the first experiment, 18 in the control experiment and 19 in the second experiment. In order to find out whether first-order and metacognitive performance differed we calculated first-order task sensitivity (because the data fell apart in three conditions we calculated d_a_, see Macmillan and Creelman, 2004), metacognitive sensitivity (meta-d_a_) and metacognitive efficiency (meta d_a_ – da, see Maniscalco and Lau, 2012; Fleming and Lau, 2014), for each condition separately. First-order task sensitivity (d_a_) and metacognitive sensitivity (meta-d_a_) are bias-free measures of the ability to distinguish two signals from each other and the ability to distinguish between correct and incorrect decisions, respectively (both in units of first-order d_a_). Metacognitive efficiency reflects metacognitive sensitivity relative to different levels of first-order task performance, which is important because metacognitive sensitivity is known to be influenced by first-order task performance (Fleming and Lau, 2014).

We performed three repeated measures analyses of variance (ANOVA) on first- and second-order task performance (d_a_, meta-d_a_, and metacognitive efficiency) with condition as the independent variable. All behavioral analyses were performed using JASP (Version 0.8.3.1), Matlab (Matlab 12.1, The MathWorks Inc.), type 2 SDT scripts (Maniscalco and Lau, 2012) and SPSS (IBM SPSS Statistics, 22.0). For the Bayesian analysis in JASP a Cauchy prior distribution centered around zero was used with an interquartile range of r = 0.707.

### EEG measurements and analyses

EEG was recorded and sampled at 1048 Hz using a Biosemi ActiveTwo 64-channel system, with four additional electrodes for horizontal and vertical eye-movements, each referenced to their counterpart (Biosemi – Amsterdam, The Netherlands). High-pass filtering (0.5 HZ), additional low-pass filtering (100 HZ) and a notch filter (50 HZ) were used. Next, we down-sampled to 512 Hz and corrected for eye movements on the basis of Independent Component Analysis (Vigário, 1997). The data was epoched −1.5 s to + 0.5 sec preceding confidence judgments. We removed trials containing irregularities due to EMG or other artifacts by visually inspecting all trials. To increase spatial specificity and to filter out deep sources we converted the data to spline Laplacian signals (Cohen, 2015). We used a sliding window Fourier transform (Mitra and Pesaran, 1999), window length: 400 ms, step size: 50 ms, to calculate the time-frequency representations of the EEG power (spectrograms) for each channel and each trial. We used a single Hanning taper for the frequency range 3–30 Hz (frequency resolution: 2.5 Hz, bin size: 1 Hz [Kloosterman et al. 2015]). To examine the way information might be distributed during metacognitive decision-making, we assessed measures of interregional functional connectivity in the beta range. In our previous study, we specifically observed effects in prefrontal channel Fz related to metacognitive performance (Wokke et al. 2017). Therefore, we specifically examined consistencies of the difference of time–frequency phase values between motor channels (C3/C4, depending on the hand that responded) and central frontal electrode Fz (Intersite Phase Clustering (ISPC), see Siegel et al. 2012; Cohen 2014) in the 500ms time period immediately preceding the confidence judgment (see Figure 1). We used ISPC measurements to determine whether reducing the amount of motor information available at the moment of confidence judgments changed the level of functional connectivity (i.e., alpha/beta phase synchronisation) between central prefrontal (Wokke et al. 2017) and motor regions. In experiment 2 three participants had to be excluded from further EEG analyses due failed EEG recordings.

Power modulations were characterized as the percentage of power change at a given time and frequency bin relative to baseline power value for that frequency bin. The baseline was calculated as the mean power across the pre-stimulus interval (from - 0.3 to 0 s relative to stimulus onset). All signal processing steps were performed using Brain Vision Analyzer (BrainProducts) and Matlab (Matlab 12.1, The MathWorks Inc.), **X** code (Cohen, 2014) and Fieldtrip (Oostenveld et al. 2011).

## Results

### Behavior

To determine whether action processes (i.e., response competition, ‘ease’ of action preparation [Wenke et al., 2010]) contributed to the quality of metacognitive judgments, we varied the amount of first-order action information present at the moment metacognitive decisions had to be provided (see Figure 1a). We constructed three conditions that differed in the moment participants had to provide their metacognitive judgment (see methods). In the first condition, participants provided verbal metacognitive judgments after the response cue and after the first-order response (ACT condition). In the second condition, metacognitive judgments were provided before the first-order response but after the presentation of the response cue (PRE_ACT condition). In the third condition, participants provided metacognitive judgments before presentation of the response cue and execution of the first-order response (PRE_CUE condition). We performed three repeated measures ANOVAs (the three conditions as levels) on first-order task performance (d_a_), metacognitive sensitivity (meta d_a_) and metacognitive efficiency (meta d_a_ - d_a_), respectively (see methods). Metacognitive sensitivity quantifies (in units of d_a_) how well a participant can discriminate correct from incorrect decisions on a first-order task. Metacognitive efficiency is the ability to discriminate between correct and incorrect decisions relative to different levels of first-order task performance^22^. Because of the known influence of first-order task performance on metacognitive performance (meta d_a_), metacognitive efficiency is a measure of metacognitive performance that is more independent from variability in first-order performance (Fleming and Lau 2014).

We found a significant effect of condition, specifically for metacognitive efficiency (*F_(2, 28)_* = 4.04 p = 0.0029, η^2^=0.224). For both d_a_ (*F_(2, 28)_* = 0.631 p =0.540, η^2^=0.043) and meta d_a_ (*F_(2, 28)_* = 1.882 p = 0.171, η^2^=0.118) no significant effects were observed. Next, we performed (one-tailed) t-tests to find out whether metacognitive efficiency decreased when response information was reduced. Results demonstrate that ACT and PRE_CUE significantly differed from each other *(t_(14)_*=2.45, p= 0.014, *d=* 0.663, BF_+0_=4.75), while no significant differences were observed between ACT and PRE_ACT (*t_(14)_*=1.65, p= 0.061, *d=* 0.426, BF_+0_=1.47) and PRE_ACT and PRE_CUE (*t_(14)_*=1.45, p= 0.085, *d=* 0.374, BF_+0_=1.13), see Figure 1b. These findings sugggest that participants’ capacity to distinguish accurate from inaccurate decisions increased when first-order response information was available compared to when such information was unavailable. We did not observe differences in the average confidence level between the conditions (all ts<0.753, ps>0.464).

To assess whether differences in the time between stimulus offset and response affected performance (e.g., due to prolonged evidence accumulation), we post-hoc explored differences between d_a_ scores in ACT and PRE_ACT, and in ACT and PRE_CUE respectively (see Figure 1b). We did not observe any significant d_a_ differences (ACT vs. PRE_ACT: *t_(14)_*=0.56, p= 0.584, BF_10_=0.301; ACT vs. PRE_CUE: *t_(14)_*=1.00, p= 0.334, BF_10_=0.403). These findings indicate that the presented blank after stimulus offset most likely eliminated effects of prolonged evidence accumulation.

### EEG results

In order to examine the neural mechanisms that support communication between motor areas and prefrontal regions during metacognitive decision-making, we assessed differences in interregional functional connectivity (beta phase synchrony) between the central frontal electrode Fz (see methods) and motor channels C3 or C4 (depending on the hand that responded) in the 500 ms time window preceding participants’ metacognitive judgment. There was a significant effect of condition for changes in beta phase synchrony (Greenhouse-Geisser corrected: *F*_(1.29,18.19)_= 8.434, *p*= 0.006, η^2^=0.376). Because oscillatory activity in the alpha band has also been closely linked to action mechanisms (Brinkman et al. 2014), we explored whether differences between conditions in alpha phase synchrony could be observed. No effects were found for changes in alpha phase synchrony between conditions (*F*_(2,28)_=1.483, *p*= 0.244, η^2^=0.096); see Figure 2. We found higher functional connectivity (beta phase synchrony) in ACT compared to PRE_ACT (*t_(14)_*=3.89, p= 0.002, *d=* 1.004, BF_10_=25.437) and PRE_CUE (*t_(14)_*=2.446, p= 0.028, *d=* 0.632, BF_10_=2.405). No differences were observed between PRE_ACT and PRE_CUE (*t_(14)_*=1.20, p= 0.250, *d=* 0.310, BF_10_=0.482).

**Figure 2.**
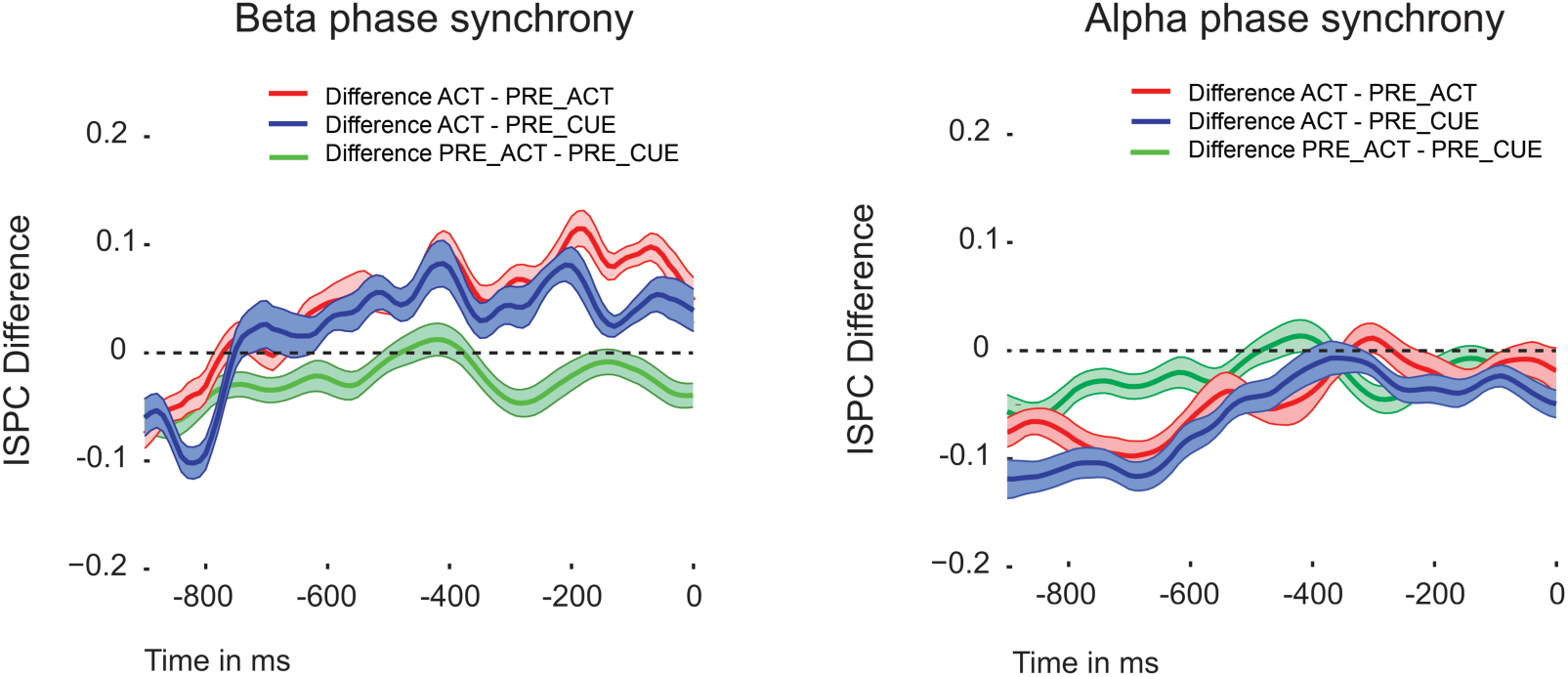
Functional connectivity. Functional connectivity (beta phase synchrony) between motor cortex and prefrontal cortex was higher in ACT where response information was available during metacognitive decision-making compared to PRE_ACT and PRE_CUE. No effects were observed for alpha phase synchrony. Shaded areas represent within-subjects standard error of the mean. Time zero refers to the onset of the metacognitive question (see Figure 1).

Next, we investigated whether functional connectivity changes (beta phase synchrony) were accompanied by changes in beta power in the central frontal channel Fz. Beta power was higher in ACT compared to PRE_ACT (*t_(14)_*=2.765, p= 0.015, *d=* 0.714, BF_10_=3.957), while no differences were found between ACT and PRE_CUE (*t_(14)_*=1.364, p= 0.194, *d=* 0.352, BF_10_=0.011); see Figure 6a.

### Control experiment

In our EEG analyses, we attempted to minimize the effect of the mere presence of a motor response (the act of moving your finger) by focusing on the last 500 ms preceding the metacognitive judgment (see Figure 1a). Nonetheless, EEG results observed in the first experiment could still be influenced by epiphenomenal/lingering motor activity caused by pressing a button in ACT versus not having pressed a button in PRE_ACT and PRE_CUE. We thus repeated the first experiment (ACT and PRE_ACT) while replacing the verbal confidence judgment with the verbal report of a random letter (see Figure 3). In this way, we were able to find out whether the observed beta effects (phase synchrony/power) were related to epiphenomenal motor activity or whether this was instead specifically linked to metacognitive judgments. In the control experiment, no differences in first-order performance (d_a_) between the two conditions were observed (*t_(18)_*=0.164, p=0.872, *d=* 0.038, BF_10_=0.240; Mean d_a_ condition 1= 0.99, SD= 0.45; Mean d_a_ condition 2= 0.97, SD= 0.44). In contrast to the first experiment, we did not observe a significant difference in functional connectivity between ACT and PRE_ACT (beta phase synchrony*: t_(18)_*=0.475, p= 0.641, *d=* 0.109, BF_10_=0.263; alpha phase synchrony: *t_(18)_*=0.511, p= 0.615, *d=* 0.117, BF_10_=0.267), see Figure 5a. Similarly to the first experiment, however, we did observe a difference in beta power between ACT and PRE_ACT (*t_(18)_*=5.098, p< 0.001, *d=* 1.201, BF_10_=311.7), see Figure 6b. These findings indicate that the increase in functional connectivity (beta phase synchrony) between frontal and motor areas is not merely caused by epiphenomenal first-order response activity, but seems instead to be connected to the metacognitive processes that follow first-order responses. In contrast, beta power differences between the conditions in the current experiments seem to be non-specific to what happens after the first-order response: we observed beta power differences when a metacognitive judgment had to be provided as well as when a random letter had to be reported.

**Figure 3.**
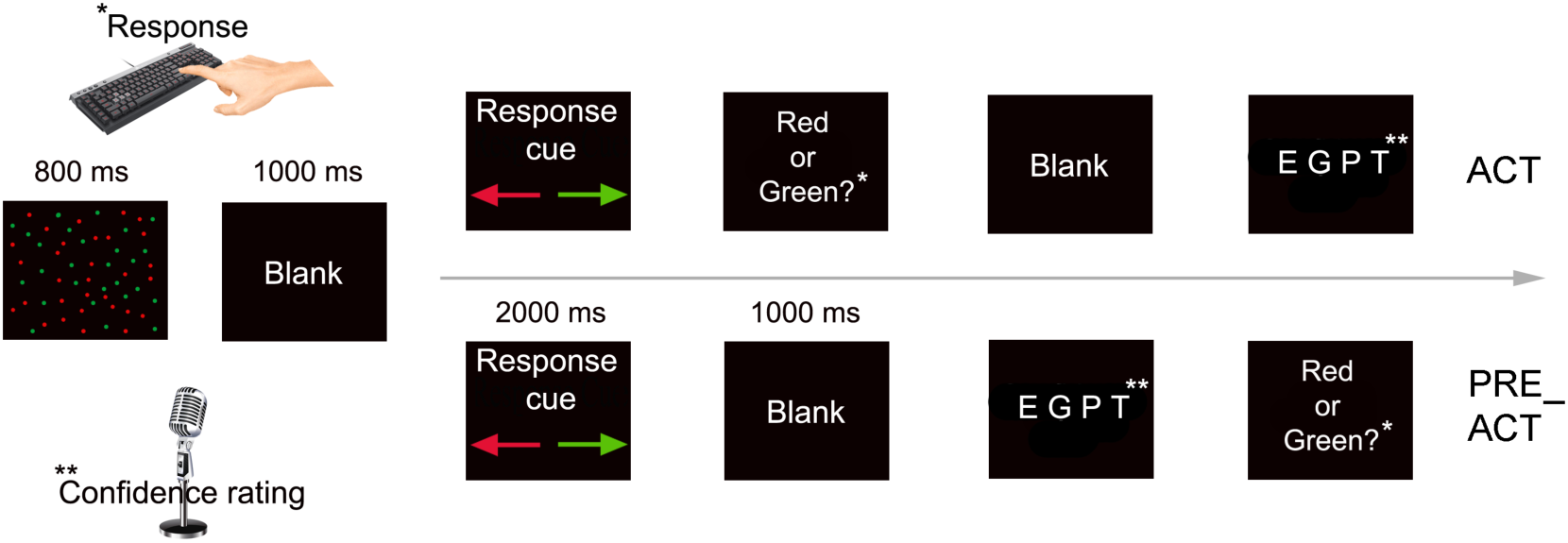
Task design control experiment. In the control experiment we replaced the metacognitive decision with a verbal response of a letter, while keeping the rest of the design identical to ACT and PRE_ACT of the first experiment.

### Experiment 2

To find out if we could replicate the findings from the first experiment and to investigate whether the strength of the stimulus-response mapping influenced the strength of the observed behavioral and EEG effects, we recorded behavioral data and EEG signals during a second experiment in which we omitted the response cue (see Figure 4a). As such, the experiment was similar to the first experiment with the exception that stimulus-response mappings were kept stable across the entire experiment.

**Figure 4.**
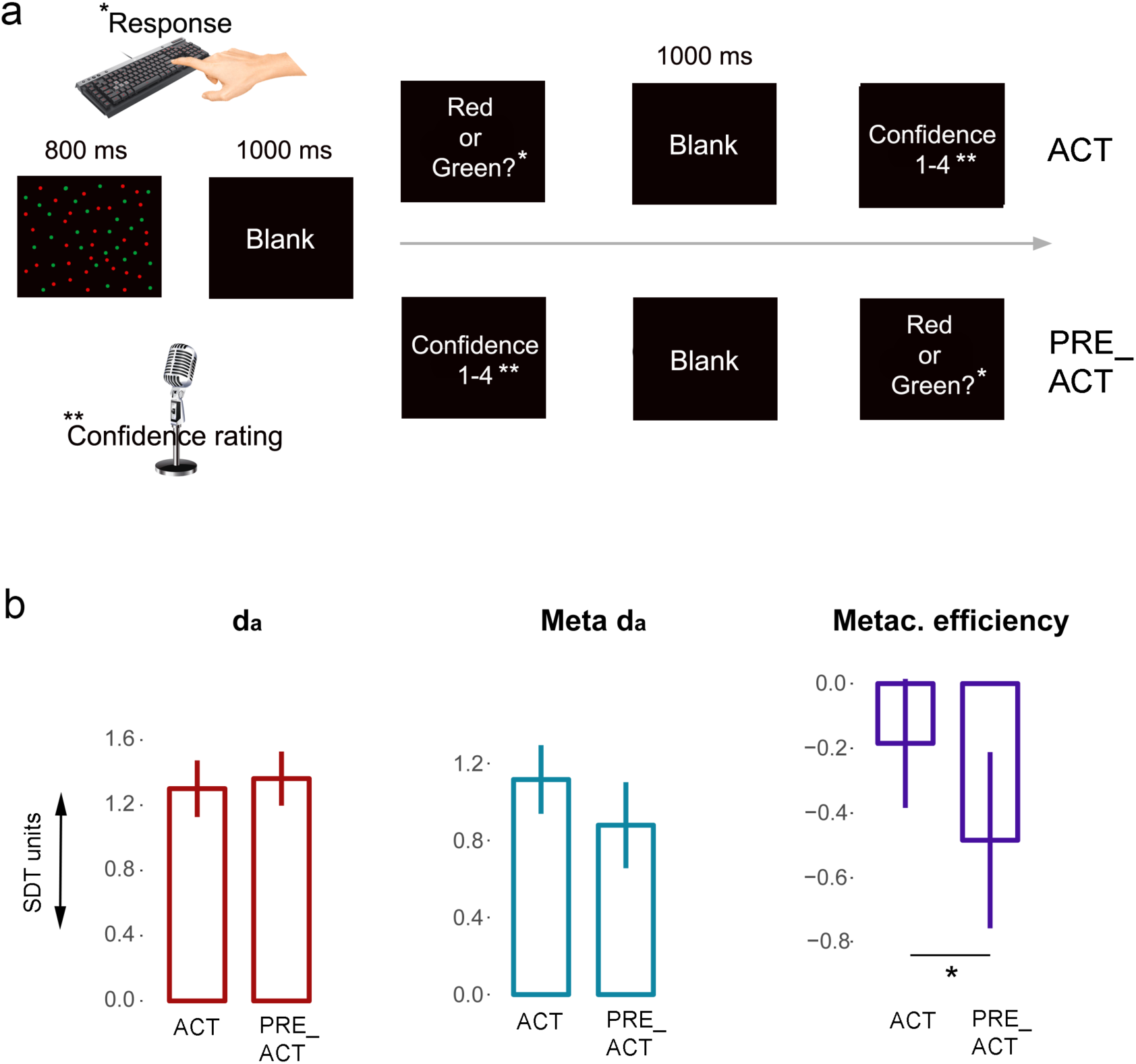
a) *Task design experiment 2.* In the second experiment we omitted the response cue, while keeping the rest of the design similar to experiment 1. b) *Behavioral results.* We replicated our findings from the first experiment and observed that metacognitive efficiency decreased when action information was absent, while first order performance remained unaffected. Error bars represent within-subjects standard error of the mean.

### Behavior

We performed (one-tailed) t-tests to find out whether metacognitive efficiency decreased when action information was absent. We replicated findings from the first experiment (though the statistical effect is rather small) and found increased metacognitive efficiency when response information was available (ACT) compared to PRE_ACT in which this information was absent *(t_(18)_*=2.134, p= 0.023, *d=* 0.490, BF_+0_=2.89). No significant differences were observed between conditions for d_a_ scores (*t_(18)_*=0.713, p= 0.758, *d=* 0.164, BF_+0_=0.151) or meta d_a_ scores (*t_(18)_*=1.622, p= 0.061, *d=* 0.372, BF_+0_=1.337), see Figure 4b. In this experiment, we did observe a consistent lower level of confidence in ACT (mean=2.63, SD=0.433) compared to PRE_ACT (mean=2.70, SD=0.431), *t_(18)_*=2.999, p= 0.012, *d=* 0.642, BF_10_=4.17.

### EEG results

In the second experiment we repeated the analyses from the first experiment by focusing on functional connectivity differences between ACT and PRE_ACT. We replicated our previous findings and observed higher functional connectivity (beta phase synchrony) in ACT compared to PRE_ACT (*t_(15)_*=4.038, p= 0.001, *d=* 1.009, BF_10_=36.003; alpha phase synchrony: *t_(15)_*=0.881, p= 0.392, *d=* 0.22, BF_10_=0.358), see Figure 5a. We also observed higher beta power in ACT compared to PRE_ACT (*t_(15)_*=2.639, p= 0.019, *d=* 0.660, BF_10_=3.269, see Figure 6c), however, due to a similar beta power effect observed in the control experiment, it is highly unlikely that the beta power effects are the result of our experimental manipulation.

**Figure 5.**
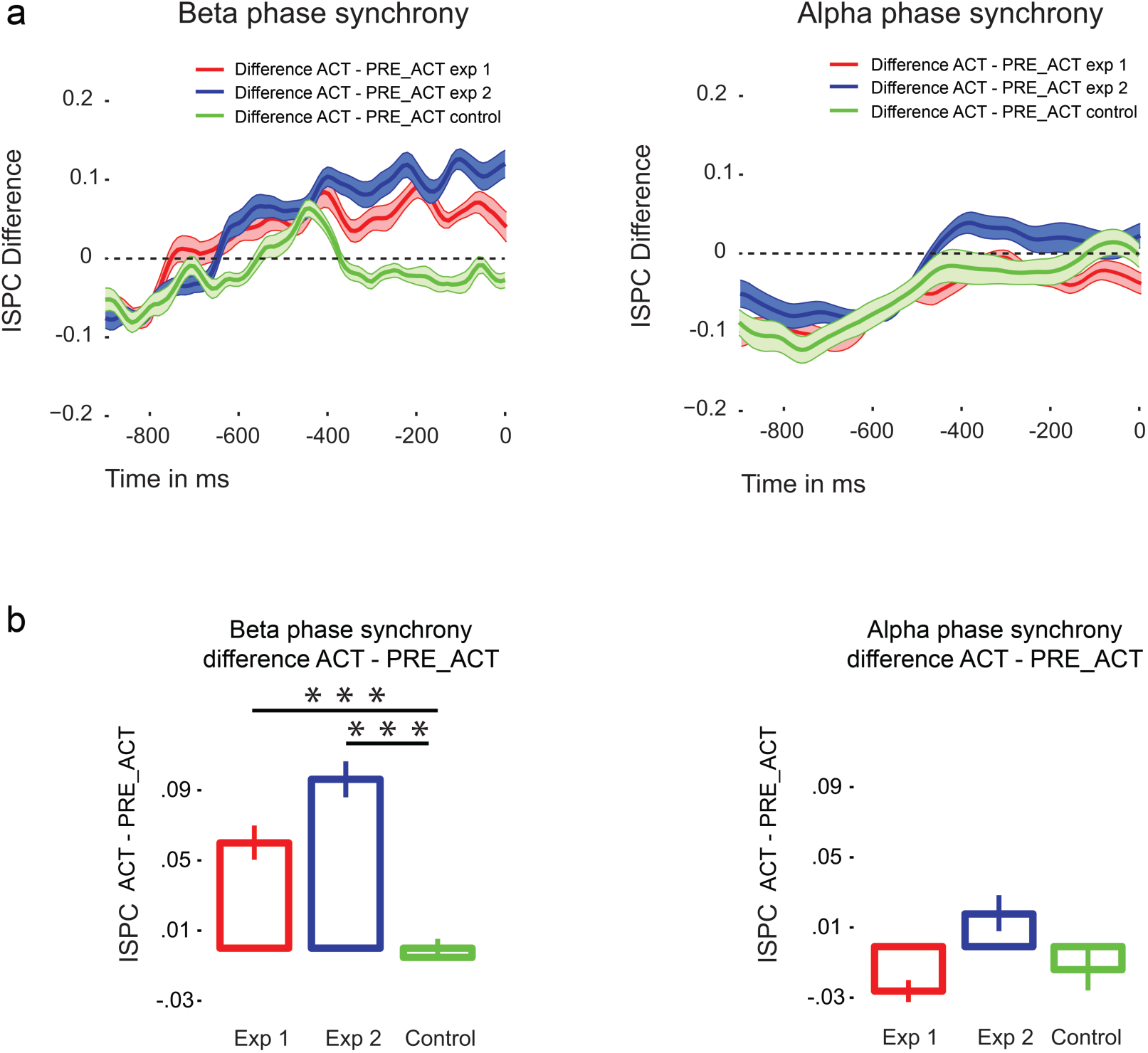
a) Functional connectivity differences of beta (left) and alpha (right) phase synchrony. Similar to Figure 2, we observed enhanced functional connectivity (beta phase synchrony) between motor cortex and central frontal cortex in ACT where response information was available during metacognitive decision-making compared to PRE_ACT. This effect was not observed in the control experiment where participants were not engaged in a metacognitive task. In all three experiments, no alpha phase synchrony differences were observed. Shaded areas represent within-subjects standard error of the mean. b) Direct comparisons of the observed beta phase synchrony differences in all three experiments show that the effect is specific to settings in which metacognitive decisions are required. Error bars represent within-subjects standard error of the mean. Time zero refers to the onset of the metacognitive question (see Figure 1).

**Figure 6.**
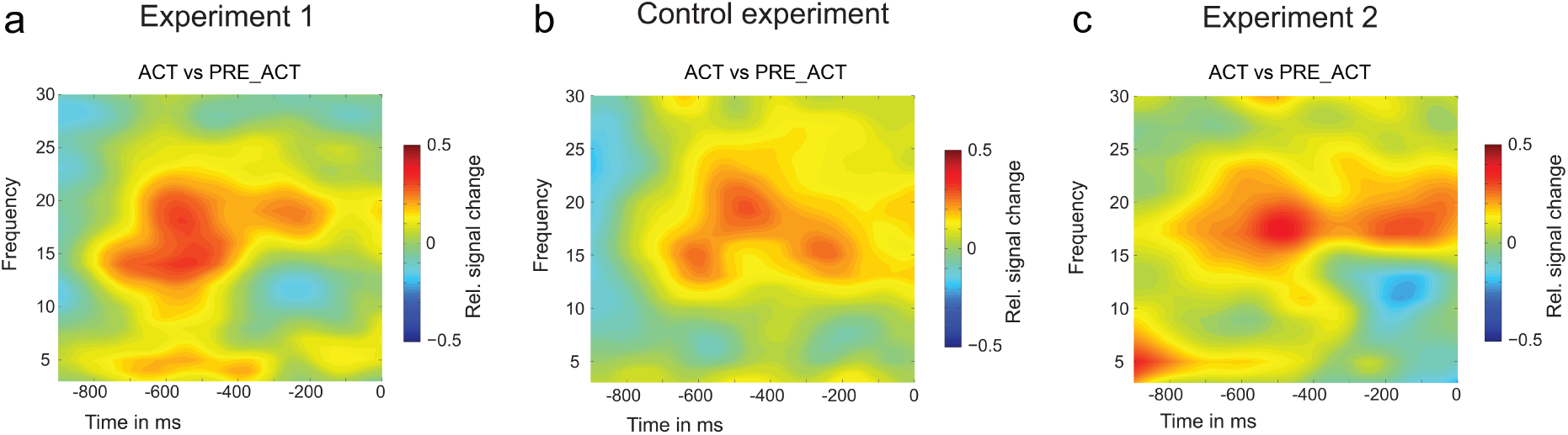
Time frequency results of experiment 1 (a), control experiment (b) and experiment 2 (c). In contrast to the functional connectivity results, we observed a similar pattern of enhanced beta power in all three experiments (including the control experiment), indicating that these beta power effects are unspecific to metacognitive decision-making. Time zero refers to the onset of the metacognitive question (see Figure 1).

### General results

In order to determine the overall effect of action processes on metacognitive efficiency, we grouped the data from the first and second experiment together (see methods) using Bayesian statistics, which make it possible to meaningfully aggregate subjects and/or experiments in a post-hoc manner. We therefore grouped PRE_ACT and PRE_CUE from experiment 1 so as to create two conditions, as in experiment 2. We observed strong evidence for higher metacognitive efficiency (BF_+0_=19.151, see Figure 7) when action information was available during metacognitive judgments. Note that the combined effect is much stronger than the weak behavioral effects observed in each individual study, suggesting the need for large enough sample size. Future studies investigating changes in metacognitive performance could benefit from such a larger sample size, and from using a staircase procedure for second-order performance as well as first-order performance, preventing the exclusion of participants.

**Figure 7.**
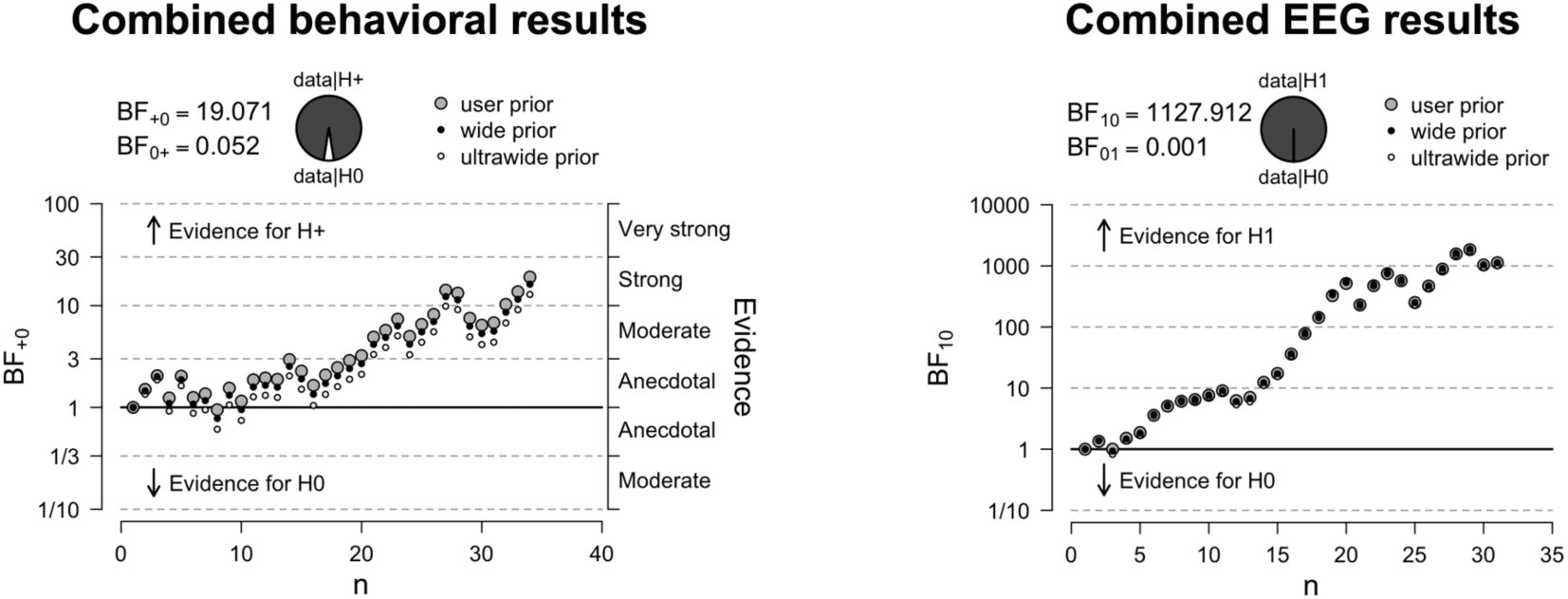
Combined results. When combining the data from experiment 1 and 2 we find strong evidence for increased metacognitive efficiency when action information is available during metacognitive decision-making. Similarly, strong evidence is observed for increased functional connectivity (beta phase synchrony) between motor channels and central frontal regions when action information is available during metacognitive decision-making.

To test whether functional connectivity differences between ACT and PRE_ACT differed between the experimental and control experiment, we directly compared ACT and PRE_ACT differences with each other (Nieuwenhuis et al. 2011) using independent sampled t-tests. In all experiments we subtracted values from PRE_ACT from ACT. Again we averaged PRE_ACT and PRE_CUE from experiment 1 and subtracted that from the ACT condition. We observed significantly greater differences in the experimental conditions compared to the control condition (first experiment vs. control experiment: *t_(32)_*=2.904, p= 0.007, *d=* 1.003, BF_10_=6.901; second experiment vs. control experiment: *t_(33)_*=4.057, p< 0.001, *d=* 1.377, BF_10_=87.51), see Figure 5b. When examining the combined data from the first and second experiment with respect to functional connectivity, we find strong evidence for greater beta phase synchrony between motor and central frontal regions when action information is available at the moment of metacognitive decision-making (BF_+0_=1127.912, see Figure 7 & 8).

**Figure 8.**
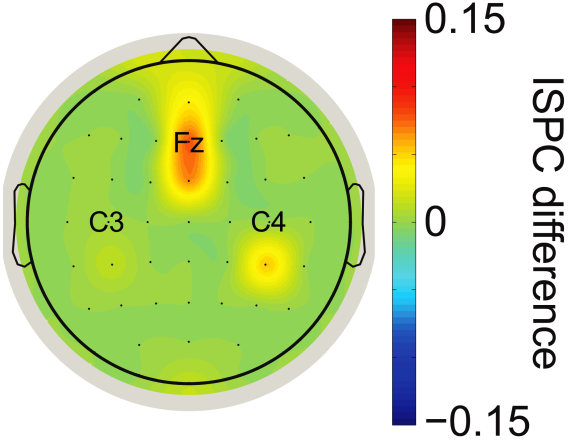
Topoplot of the combined functional connectivity effect (ACT vs. PRE_ACT). For illustration purposes we plotted beta phase synchrony differences between ‘seed’ electrode C3/C4 and other electrodes to show the spatial distribution of the observed effect.

## Discussion

Decision-making is typically accompanied by an estimate about the quality of ones choices, actions or performance. Adequate metacognition is not only important in everyday life settings (e.g., using the right amount of ingredients while cooking, or knowing what you know while studying for an exam), but can even be critical in certain situations (e.g., in case of medical decisions, or decisions made by a flight controller). Despite its importance, it remains unclear how metacognition emerges, and what kind of information is used to determine the quality of our decisions.

Here, we investigated whether first-order action information could inform second-order (metacognitive) decisions. Specifically, we studied whether reducing available first-order response information at the moment second-order decisions had to be provided affected metacognitive performance in a color discrimination task. Further, we investigated whether functional connectivity between motor regions and prefrontal cortex could serve as a mechanism to convey action information during metacognitive decision-making. Results demonstrate that metacognitive efficiency slightly decreased when first-order action information was reduced at the moment metacognitive decisions had to be provided. We replicated our findings in a second experiment and showed that the effect was small but robust to changes in the experimental design (see Figure 1b, 4b & 7). Similarly, we found converging electrophysiological evidence that functional connectivity between motor areas and prefrontal cortex increases during metacognitive decision-making when action information is available (see Figure 2 & 5). In a control experiment, we demonstrated that this effect was not related to lingering response activity, but in fact specific to metacognitive processes following first-order decisions (Figure 5). Combined analyses of the three experiments provide converging evidence for the contribution of action information in metacognitive decision-making.

### Models of metacognitive decision-making

In lab settings, metacognition is typically studied by asking participants to make a decision about a stimulus (e.g., the motion direction of a cloud of moving dots, the orientation of a grating), after which they are asked to provide the level of confidence in their decision being correct. Previously, it has been shown that manipulating stimulus parameters (evidence strength and evidence reliability) affects confidence judgments (Boldt et al. 2017) during perceptual decision-making, suggesting similar (sensory) evidence processing mechanisms support first- and second-order decision-making. Similarly, in signal-detection-like models, the distance of the decision variable from a criterion represents a level of confidence (Macmillan and Creelman 2004; Kepecs et al. 2008; Kiani and Shadlen 2009; Fetsch et al. 2014). The time between the decision and presentation of sensory evidence could in such cases result in discrepancies between first- and second-order decisions, due to prolonged accumulation of evidence (Berg et al. 2015; Boldt and Yueng 2015; Calderon et al. 2018). Alternatively, different sources or quality of information could contribute to first- and second-order decisions (Pleskac and Busemeyer 2010; Charles et al. 2014), resulting in different first- and second-order performance. With respect to the latter, we previously demonstrated that sensory evidence contributing to first-order decision-making does not similarly support metacognitive decision-making. Variance in first-order performance was driven by different stimulus features compared to variance in metacognitive performance. These findings indicated that sensory evidence used for first-order performance differed from information used for metacognitive judgments (Wokke et al. 2017). Maniscalco and Lau recently compared models describing discrepancies between first- and second-order decisions during a visual masking task (Maniscalco and Lau 2016). They compared models which depict first- and second-order decision-making as supported by similar sources of information (single channels models) with dual channel models, which describe two processing streams giving rise to first- and second-order task performance; and hierarchical models, which presume that a late processing stage monitors the state of sensory processing. Their results demonstrated that dissociations between first- and second-order performance are best captured by hierarchical models. However, in their study they used data from a visual masking task using visibility ratings as a second-order task. It therefore remains unclear how their results generalize to different tasks and metacognitive measures. Fleming and Daw (2017) recently put forward a framework in which confidence operates as a second-order computation about one’s own performance. While first-order models are able to reproduce the above-described relationship of confidence and stimulus parameters, their second-order model accommodates the present findings that action information influences metacognitive performance and metacognitive bias. The second-order framework predicts that action affects confidence ratings, in the sense that it decreases overall confidence and enhances metacognitive performance. In the current experiments we observed this pattern in our behavioral results. In two experiments, we demonstrated that metacognitive efficiency increased when first-order action information became available for second-order decision-making. In addition, we observed a (somewhat counterintuitive) decrease in confidence when metacognitive judgments followed first-order responses in the second experiment, as predicted by the second-order model (Fleming & Daw, 2017). We did not observe differences in overall confidence in the first experiment. It could be that trial-by-trial alternations of stimulus-response mappings in the first experiment tampered the effect on metacognitive bias shifts. Previously, it was found that participants’ metacognitive bias shifted when they learned motor sequences in a blocked design compared to when sequences were interleaved (Simon and Bjork 2001). These findings suggest that the current ease of stimulus-response mappings affected metacognitive bias. In that sense, it would be interesting for future experiments to assess whether/how manipulation of ease or the integrity of first-order responses influences metacognitive behavior.

### Beta oscillations

Beta oscillations are classically linked to sensory and motor processing (Pfurtscheller and Lopes da Silva 1999; Spitzer and Haegens 2017). During preparation and execution of movements, beta band activity typically decreases initially, followed by an increase in beta power (Kilavik et al. 2013). For instance, an upcoming action could be reliably predicted several seconds prior to response execution, based on lateralization of beta band activity in motor regions, linking beta band activity to the unfolding of an action (Donner et al 2009). It has been suggested that beta activity reflects the maintenance of an existing motor set whilst weakening processing of new actions (Gilbertson et al. 2005). Interestingly, beta synchronization has been associated with the correctness of an action and has been shown to follow motor errors or after observing the motor errors of others (Koelewijn et al. 2008; Swann et al 2009). Recently, the importance of beta oscillations has been demonstrated beyond the sensorimotor domain, extending to visual perception (Piantoni et al. 2010; Kloosterman et al. 2015; Bastos et al. 2015), working memory (Siegel et al. 2009), long-term memory (Hanslmayr et al. 2016), and decision-making (Donner et al. 2009; Haegens et al. 2011; Wyart et al. 2015; Herding et al. 2016). It has been proposed that beta oscillations support long-range neuronal interactions (Thompson and Varela 2001; Siegel et al. 2011; Benchenane et al. 2011) thereby maintaining a current cognitive set, sensorimotor state or the so-called ‘status quo’ (Engel and Fries 2010). In this way, the up or down regulation of beta depends on whether the ‘status quo’ is prioritized over novel incoming signals. Recently, Haegens and Spitzer (2017) extended the role of beta oscillations further, advocating a role of beta in the awakening of a (endogenous) cognitive set, depending on current task demands.

In the current study, we found increased phase synchrony in the beta band between motor channels and central frontal regions (electrode Fz) specifically when a metacognitive decision followed the first-order response. Critically, when task demands changed and a metacognitive judgment was not required, beta phase synchrony differences between conditions disappeared. In line with the above-proposed role of beta oscillations, our beta phase synchrony findings indicate that task demands (the metacognitive task) resulted in the maintenance of first-order action information (e.g., response fluency, response competition strength). It would be interesting to investigate what role explicitly asking for a metacognitive judgment has on beta band activity. If we assume that decisions are naturally accompanied by an estimate about the quality of an action or choice, it could be that by explicitly asking for such an estimate after a short time interval we could have prolonged or boosted a naturally occurring more transient event (for a similar discussion in consciousness research see: Tsuchiya et al. 2015). Indeed, beta phase synchrony effects in the control condition initially seem to mimic those observed in the other two experiments, only starting to deflect in the period preceding the metacognitive judgment. It would be interesting to test ‘naturally occurring’ metacognitive processes in future experiments, thereby using observed neural markers of explicitly probed metacognitive processes (Fleming and Dolan 2012; Fleming et al. 2012; Murphy et al. 2015; Wokke et al. 2017).

### Motor activity and metacognition

The present results indicate a contribution of first-order motor response information in metacognitive decision-making. Previously, Wenke and colleagues (2010) demonstrated that participants were sensitive to conflicting motor activity (response competition) induced by subliminal information. In their study the ‘‘ease” or “smoothness” of action selection in a visual reaction-time task was manipulated by presenting a subliminal response prime that was congruent to one out of two action possibilities. Results demonstrated that action priming influenced the sense of control over action consequences following the response. Other work indicates that metacognitive experience of response competition is crucial for triggering cognitive adaptation (Desender et al. 2014; Questienne et al. 2016). Further, it has been shown in the memory domain that the experience of motor fluency is used as a cue that affects metamemory (Susser and Mulligan 2015; Susser et al. 2017).

Recently, it has been shown that perceptual decisions were biased by the amount of motor effort it took for participants to make the response (Hagura et al. 2017). In this study, participants’ decision was biased towards the least effortful motor response. These findings demonstrate that the ease to act on a decision might influence the decision itself. However, it seems that metacognitive awareness of effort or of task demands is necessary for the development of such a decision bias (Desender et al. 2017). In the current experiments, results show that participants could be sensitive to response competition, the fluency or ease of the first-order response (Pacherie 2008; Questienne et al. 2016) when computing an estimate about the quality of the decision.

Alternatively, motor activity could provide insight into the mechanisms of the unfolding perceptual decision. Recent studies demonstrated that evidence accumulation processes ‘echo’ in activity in motor regions (Donner et al. 2009). As such, perceptual and cognitive states could be reflected in the motor system (Song and Nakayama 2009; De Lange et al. 2013) and be used to inform metacognitive decisions.

### Prefrontal cortex and metacognition

Previous work demonstrated that lesions to prefrontal cortex affect metacognitive performance without altering first-order decision-making (Pannu and Kaszniak 2005; Fleming et al. 2014). Similarly, disrupting prefrontal activity via theta burst stimulation has been shown to selectively alter metacognitive performance (Rounis et al. 2010; Ryals et al. 2016; Shekhar and Rahnev 2018, but see Bor at al. 2017 and Ruby et al. 2018). The detection of erroneous behavior, a key aspect of metacognition (Yueng and Summerfield 2012; Fleming and Daw 2017), has been strongly linked to a rapidly emerging central frontal negativity in the EEG signal (error-related negativity, Falkenstein et al. 1991), thought to reflect coordinated theta oscillatory mechanisms (Luu and Tucker 2001; Cohen et al. 2008; Bates et al. 2009; Cavenagh et al. 2009; Cohen and Cavenagh 2011). In addition, theta has been implicated in learning, feedback processing, and action monitoring (Jensen and Lisman 2000; Dragoi and Buzsáki 2006; Sauseng et al. 2006; Cohen and Cavenagh 2011; Van de Vijver et al. 2011; Cavenagh and Frank 2014; Van Driel et al. 2015). Recently, fluctuations in prefrontal theta band activity has been linked to fluctuations in metacognitive performance (Murphy et al. 2015; Fleming 2016; Wokke et al. 2017). Taken together, these findings suggest that frequent exposure to external feedback, learning from ones correct and incorrect decisions induces a shift in which error detection, initially elicited by external feedback, is shifting towards the use of internal representations of learned stimulus-response contingencies. This internally processing of the probabilities of our actions towards outside events and their most likely outcomes (Holroyd and Coles 2002; Cleeremans et al. 2007; Cleeremans 2011; Yeung and Summerfield 2012) could be used to adapt future behavior. In such a way, metacognition could be seen as an internalization of external feedback processing and error monitoring, employing similar neural mechanisms (Buzsáki et al. 2014; Murphy et al. 2015).

It has been previously proposed that next to perceptual evidence, inferences about “the state of the decider” (i.e., one’s own actions [Fleming and Daw, 2017], and prior or global estimates of performance [Roualt et al. 2019]) are important for metacognitive decision-making. In addition, to adequately compute an estimate about the quality of a decision it is necessary to know the broader task context or infer “the state of the world” (i.e., value for an action at a certain state of the (task) environment) at the moment of the decision (Wilson et al. 2014; Schuck et al. 2018; Schuck et al. 2016; Wokke et al. 2016). Recently, the orbitofrontal cortex has been linked to inferring such “states of the world” during decision-making (Schuck et al. 2018; Wokke and Ro 2019). As such, central frontal regions and anterior frontal areas could play distinct roles in metacognitive decision-making (see also Shekhar and Rahnev 2018). Figure 9 illustrates how sensory, action and interoceptive signals could be integrated in central frontal regions, interacting with anterior prefrontal regions providing inferences about the state of the world (Schuck et al. 2018; Wilson et al. 2014) and the state of the decider (Roualt et al. 2019; Fleming and Daw 2017) when computing an estimate about the quality of a decision.

**Figure 9.**
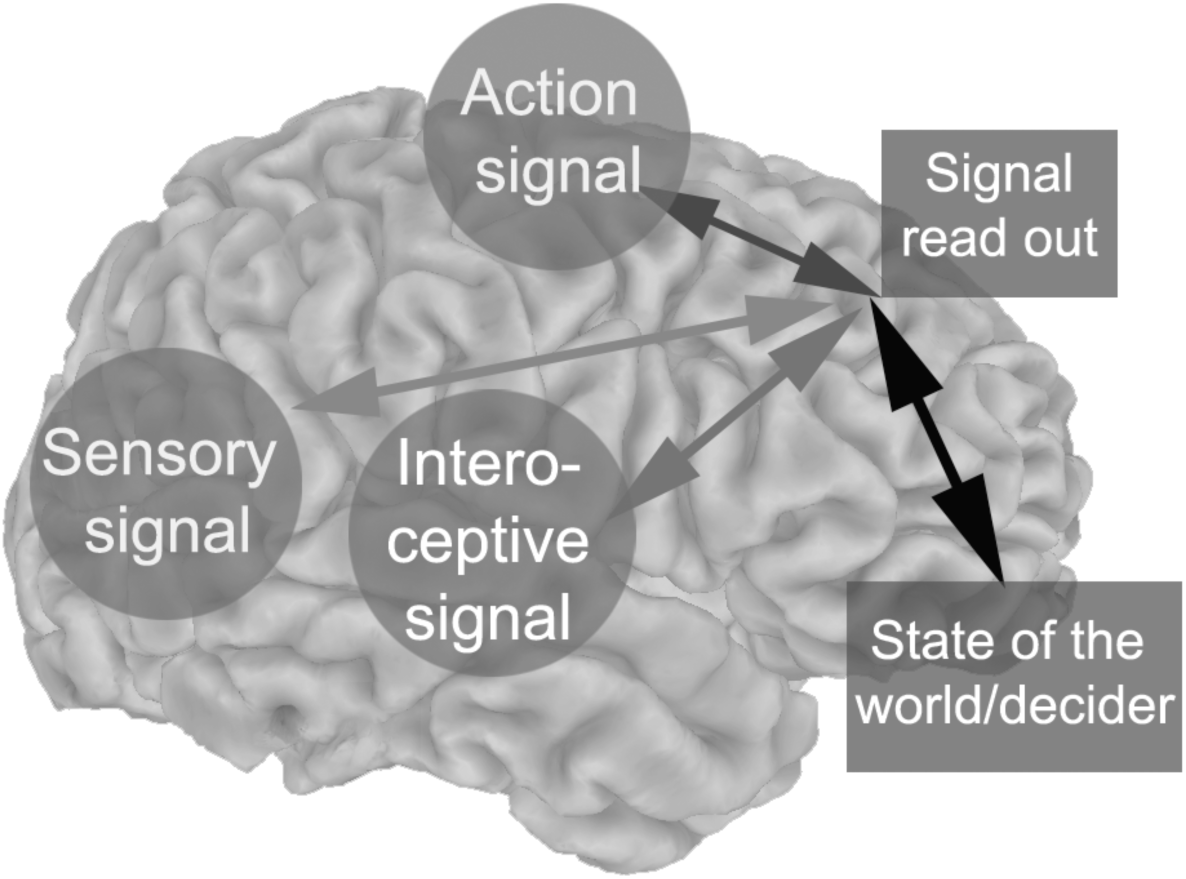
Sensory, interoceptive and action signals are read out in central frontal cortex. Anterior prefrontal cortex provides predictions about the “state of the world” and the “state of the decider” when a decision is made. Central frontal theta oscillations serve as a mechanism to broadcast the need for control in response to the estimate about the quality of the decision.

### Limitations

In the current study, we focused on functional connectivity changes between motor and prefrontal regions. However, the current neural measurements (EEG) lack spatial specificity to make strong claims about neural sources. It would therefore be necessary to replicate our findings using alternative methods (e.g. fMRI) that have a higher spatial resolution. We used a staircase performance prior to the experimental blocks to determine appropriate task settings. Despite our efforts we had to exclude participants based on first- and second-order task performance. In future studies it might be useful to use a staircase for second-order task performance in addition to first-order performance.

### Conclusion

Monitoring and evaluation of ones own performance is crucial for adept behavior. However, how metacognition emerges is still hotly debated (Berg et al. 2016; Maniscalco and Lau 2016; Fleming and Daw 2017). In a series of three experiments, we demonstrated that manipulations of available action information affected metacognitive performance. Concurrent EEG recordings showed that functional connectivity between prefrontal regions and motor areas increased after a first-order response, specifically when a metacognitive judgment was required. Together with previous findings (Fleming et al. 2015; Siedlecka et al. 2016; Fleming 2016; Wokke et al. 2017), our results demonstrate that post-decisional action information contributes to metacognitive decision-making, thereby painting a picture of metacognition as a second-order process employing endogenous control mechanisms.

## Acknowledgements

We thank Lisa Padding and Adeline de Cia for helping with data collection and Maries E. Vissers for her useful comments. This work was supported by the European Research Council (Advanced Grant RADICAL to A.C.), the European Union’s Horizon 2020 research and innovation programme under the Marie Skłodowska-Curie grant agreement (Meta_Mind - DLV-704361 to M.E.W.), and by the Belgian Science Policy Office (Interuniversity Poles of Attraction Grant P7/33 to A.C.). A.C. is a research director of the National Fund for Scientific Research and a senior fellow of the Canadian Institute for Advanced Research (CIFAR).

## Author contributions

M.E.W. and D.A. designed the study, D.A. conducted the experiment, M.E.W and D.A. analyzed the data, and M.E.W., D.A. and A.C. wrote the manuscript.

